# Loss of tRNA uridine thiolation affects mRNA translation, protein production, and sulfur-compound metabolism in Arabidopsis

**DOI:** 10.1101/2025.09.25.678457

**Authors:** Arnaud Dannfald, Adrien Cadoudal, Marie-Christine Carpentier, Rémy Merret, Quentin Rigal, Jérome Vialaret, Aurore Attina, Jana Kindermans, Christina Berrissou, Anna Koprivova, Alexandre David, Christophe Hirtz, Jean-Jacques Favory, Stanislav Kopriva, Laurence Drouard, Jean-Marc Deragon

## Abstract

The uridine at position 34 of tRNA anticodon loops is always modified at variable levels depending on environmental conditions, but a function for this highly conserved modification has not been firmly established. Using *Arabidopsis thaliana*, we show that the thiolation of U_34_ is a prerequisite for the subsequent modifications at position 32 and 37 of the tRNA^Lys^(UUU) anticodon loop, revealing a novel modification network. Surprisingly, the level of tRNA^Lys^(UUU) is strongly increased rather than reduced in the *ctu1* or *ctu2* mutant backgrounds that prevent these modifications. This suggests the existence of a regulatory feedback loop that drives the transcription of this specific tRNA gene family. Furthermore, we observed that the ability of the ribosome to decode AAA, GAA and CAA codons is impaired when the thiol group is lost, leading to a reduction in protein production, especially for genes enriched in these codons. Finally, we show that loss of tRNA thiolation results in variations in levels of many proteins involved in sulfur-compound metabolism and several sulfur-containing metabolites, suggesting that the level of tRNA thiolation may act as a sensor that regulates these processes.

## Introduction

Living organisms must differentially regulate gene expression during development or in response to environmental cues. Understanding the different underlying mechanisms responsible for key genome reprogramming events is critical, especially in the context of global warming. Modification of the genetic program can occur at different steps of gene expression, the most studied being mRNA production and post-translational protein modifications. However, mRNA post-transcriptional modifications and translation are two additional regulatory steps that are essential to achieve proper gene expression. Recently, chemical modification of mRNAs (i.e., epitranscriptomic modifications) has been identify as an important regulatory mechanism affecting mRNA maturation, stability, export and translation potential^1^. An emerging theme that has not yet been fully explored is the possibility that the translation machinery itself could adapt and more efficiently target and translate key mRNAs that contribute to genome reprogramming events^2–4^. One way to achieve this is to create so-called "specialized" ribosomes by modifying the composition of ribosomal RNA and/or proteins or the way these two types of molecules are chemically modified^3–6^. Another way would be to change the epitranscriptomic status of tRNAs, the other main partner of translation^2^. Initially tRNA epitranscriptomic marks were thought to be essential but static chemical decorations, but several reports have now shown that they can vary according to environmental conditions and propose that these variations may favor the translation of relevant mRNA populations and thus contribute to adaptation^7–16^.

Among the highly diverse chemical tags present on tRNAs^17^, the modification at position U_34_ of the anticodon loop is one of the most conserved ones, being present in bacteria, archaea and eukaryotic tRNAs^17–19^. The exact nature of the modification at this position can vary between species and between different tRNAs within a given species^17^. In eukaryotes, cytosolic tRNAs mainly contain 5-carbamoylmethyluridine (ncm^5^U), 5-methoxycarbonylmethyluridine (mcm^5^U) or 5-methoxycarbonyl-2-thiouridine (mcm^5^s^2^U) at position U_34_, although other minor forms may exist^17,20,21^. These marks are the result of the intervention of several enzymatic actors^22–24^. The elongator complex, composed of six subunits (ELP1-ELP6), and its co-factor KTI12/DRL1^25^ are required to convert U_34_ into a first intermediate (5-carboxymethyluridine, cm^5^U), followed by the TRM9/TRM112 complex, which produces mcm^5^U. Thiolation of this mark is achieved by the CTU1/CTU2 holoenzyme, with sulfur being transferred from cysteine via the Ubiquitin Related Modifier 1 (URM1) pathway^26,27^. The enzymatic activity involved in the formation of ncm^5^U from cm^5^U is currently unknown. Complete loss of the U_34_ modification is embryo lethal in *Drosophila melanogaster* and mouse^28,29^, causes severe developmental defects in nematodes (*Caenorhabditis elegans*) and plants^30,31^, and generates stress-hypersensitive phenotypes in yeast (*Saccharomyces cerevisiae*)^14,32^. A more specific loss of the thiolation mark is associated with thermotolerance defects in yeast, plants and nematodes^24,33,34^ and leads to drought hypersensitivity and root deficient phenotypes in plants^34–38^. Interestingly, the U_34_ position is usually not fully modified under normal growth conditions, and its stoichiometry varies with environmental conditions^7–10,39^. For example, thiolation of yeast tRNA is decreased under sulfur-deficient conditions, which slows growth and stimulates the synthesis and salvage of methionine and cysteine^39^.

Despite the importance of the U_34_ modification, little is known about its precise molecular impact on ribosome decoding capacity and protein production and its contribution to an appropriate response under a given physiological condition. In yeast and *Schizosaccharomyces pombe*, variations in U_34_ modifications have been shown to affect the translation of mRNAs enriched in their cognate codons, in some cases leading to a better physiological response to stress situations^14,40,41^. However, this view has been challenged by reports proposing that having a proper level of U_34_ modification is simply a way to preserve translation elongation potential and to prevent ribosome pausing and the induction of proteotoxic stress^26,32,42,43^. In this scenario, most phenotypes caused by a reduction in the level of U_34_ modification would result from a systemic failure of protein homeostasis, rather than from an inability to produce the correct amount of protein from specific mRNAs enriched in codons that depends on this modification. A crosstalk between the Target of Rapamycin (TOR) pathway and variation of U_34_ modification levels has been also proposed in yeast, human and plants^38,44,45^, but whether such variation may affect translation through the TOR pathway is still to be demonstrated. Therefore, further evidence is needed to firmly conclude that modifying the level of tRNA U_34_ chemical tags is a way to regulate gene expression by adjusting translation and protein production in response to physiological conditions. Also, little is known on the impact of U_34_ modification on tRNA stability, on the deposition of other chemical tags, especially in the anticodon loop, and how this could impact protein production and essential metabolic processes.

In this work, using *Arabidopsis thaliana* as a model, we show that tRNAs^Lys^(UUU) and tRNA^Gln^(UUG), although CTU1/CTU2 substrates, have different chemical tags in their anticodon loops. We observed that the modification of U_34_ is essential to modify tRNA^Lys^(UUU) at positions 32 and 37 and that, surprisingly, in the absence of this modification the steady-state level of this tRNA is not reduced but strongly increased. We also show that the ability of the ribosome to decode corresponding codons is impaired in the *ctu2* mutant, leading to a reduction in protein production mainly for genes enriched in these codons. Finally, we show that many proteins misregulated in *ctu2* are involved in sulfur-compound metabolism and that the concentration of key sulfur-containing metabolites (sulfate, cysteine, and glucosinolates) is affected in the mutant, suggesting that the level of tRNA thiolation works as a sensor regulating these processes.

## Results

### Analysis of the nature of chemical modifications present at the U_34_ position in Arabidopsis tRNAs: Impact on other modifications and on the tRNA global steady-state level

The uridine at position 34 of the tRNA anticodon loop is always the target of chemical modifications^17–19^, although the exact nature of these complex and highly variable modifications is often not well known for individual tRNAs. In eukaryotes, the CTU1/CTU2 holoenzyme is responsible for the thiolation step leading to the formation of mcm^5^s^2^U_34_. This modification has been found in plant cytosolic tRNA^Lys^(UUU), tRNA^Glu^(UUC) and tRNA^Gln^(UUG)^35^. To better characterize the U_34_ modification pathways in plants, we first purified total tRNAs from 15-day-old Arabidopsis WT (Col0) seedlings and from knockout mutants deficient in the production of DRL1 (also named KTI12, an essential cofactor of the ELP1-6 elongator complex^46^), TRM9, ALKBH8, CTU1, CTU2, and from a ctu2:pUBQ-CTU2 complemented transgenic line (*ctu2*-L5) (Figure 1A, Table S1). Note that in contrast to the situation in mammals, where the enzyme named ALKBH8 combines both methylase and hydroxylase activities, in Arabidopsis these two activities are provided by two different proteins, TRM9 which contains the methylase and ALKBH8 responsible for the hydroxylase part. We observed that WT Arabidopsis tRNAs mainly possess ncm^5^U, (S)-mchm^5^U, mcm^5^s^2^U at position 34 with a very small amount of mcm^5^U and cm^5^U (Figure 1A), confirming a previous report^21^. As expected, we observed a complete loss of mcm^5^s^2^U and a strong increase of mcm^5^U in *ctu1* or *ctu2*, while the levels of (S)-mchm^5^U and ncm^5^U were unaffected (Figure 1A). The WT situation is restored in the *ctu2-*L5 complemented line for all chemical marks (Figure 1A). The inhibition of the elongator complex in the *drl1* mutant prevents any type of modification at the U_34_ position. The loss of TRM9 results in strong cm^5^U accumulation and ncm^5^s^2^U detection, with no effect on ncm^5^U. It also prevents the accumulation of other modifications. Finally, mutation of ALKBH8 results in the complete loss of (S)-mchm^5^U, a strong increase of mcm^5^U and mcm^5^Um and a small increase in mcm^5^s^2^U. Based on these results, we propose a representation of the plant U_34_ modification pathway in Arabidopsis in Figure 1B.

**Figure 1.**
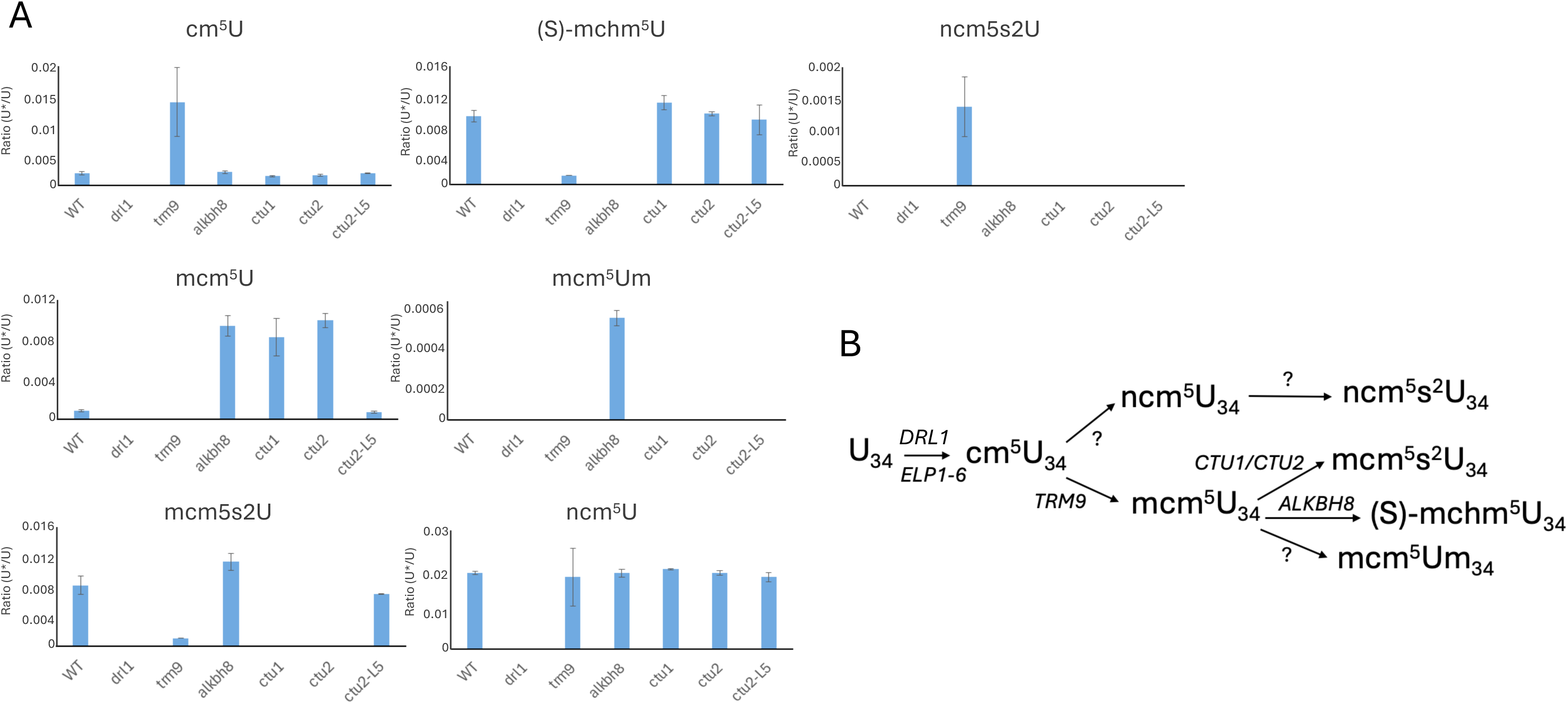
Chemical modifications present at the U_34_ position of Arabidopsis tRNAs. A) U_34_ chemical tags present in WT Arabidopsis total tRNA extracts compared to *drl1*, *trm9*, *alkbh8*, *ctu1*, *ctu2* mutant backgrounds and in a *ctu2*:pUBQ-CTU2 complemented transgenic line (*ctu2-L5*). The y-axis of the histogram represents the ratio of the area under the curve (AUC) measured by liquid chromatography/mass spectrometry (LC-MS/MS) of the modified nucleoside (U*) relative to the corresponding unmodified nucleoside (U) (mean of three biological replicates). Error bars represent the standard deviation (SD) of three biological replicates. Raw data can be found in Table S1. B) Major U_34_ modification pathways proposed for Arabidopsis cytosolic tRNAs. The first step involves the elongator complex (ELP1-6) and its co-factor DRL1 to generate the cm^5^U modifications. This mark can be modified by TRM9 to produce mcm^5^U, which can be either hydroxylated by ALKBH8 to produce (S)-mchm^5^U or thiolated by the CTU1/CTU2 holoenzyme to produce mcm^5^s^2^U. In addition, the absence of ALKBH8 or TRM9 allows for the respective detection of mcm^5^Um and ncm^5^s^2^U.

Next, we wanted to test whether different U_34_ chemical tags could coexist on a given tRNA species and whether this modification is a prerequisite for the deposition of other chemical tags in the anticodon loop. To answer these questions, we purified two of the three tRNAs that are substrates of the CTU1/CTU2 holoenzyme and known to possess mcm^5^s^2^U at position 34, tRNA^Lys^(UUU) and tRNA^Gln^(UUG), and analyzed the nature of the chemical modifications present on them by mass spectrometry (Figure 2, Table S1) (the list of chemical tags searched in this analysis is presented in Table S2). Analysis of purified tRNA^Lys^(UUU) confirmed that mcm^5^s^2^U is the only tag present at position 34 on this tRNA in WT plants (Figure 2A). In the *ctu1* or *ctu2* backgrounds, this tag is replaced by mcm^5^U and, in *trm9*, by a combination of cm^5^U, ncm^5^U and ncm^5^s^2^U. Interestingly, 2’-O-methylated cytosine (Cm), possibly at position 32, and ms^2^t^6^A, probably at position 37, were also detected. However, both modifications were drastically reduced in the *ctu1*, *ctu2* or *trm9* background, indicating that thiolation of position U_34_ is a prerequisite for the modification at positions 32 and 37 of this tRNA. Analysis of purified tRNA^Gln^(UUG) revealed that both mcm^5^s^2^U and (S)-mchm^5^U were present at position 34 in WT plants (Figure 2B), whereas this later tag was not detected in tRNA^Lys^(UUU) (Figure 2A). In the *ctu1* or *ctu2* backgrounds, mcm^5^s^2^U was replaced by mcm^5^U, while (S)-mchm^5^U was still detected but at a lower level. In *trm9*, a combination of cm^5^U, ncm^5^U and ncm^5^s^2^U was present at position 34. As recently reported^47^, we confirmed the presence of m^2^A (and not ms^2^t^6^A) at position 37 of this tRNA, while the cytosine at position 32 is unlikely to be modified (Figure 2B). The level of m^2^A was reduced in *ctu1* or *ctu2*, suggesting that U_34_ thiolation is required for full m^2^A modification of this tRNA. Also, m^2^A was almost completely lost by the transition from mcm^5^U to ncm^5^U in the *trm9* background. For both tRNAs, the m^1^A_58_ modification was detected and was not affected by the different mutant backgrounds (Figure 2A-B). Other modifications detected in this analysis that are outside the anticodon loop are shown in Figure S1. Although some marks are shared between these two tRNAs (m^1^A (see Figure 2) D, m^5^U, m^5^C, Ψ), clear differences could be observed for several marks: m^2^_2_G, m^7^G and m^1^G detected only in tRNA^Lys^(UUU) and Gm, m^2^G and Um detected only in tRNA^Gln^(UUG).

**Figure 2.**
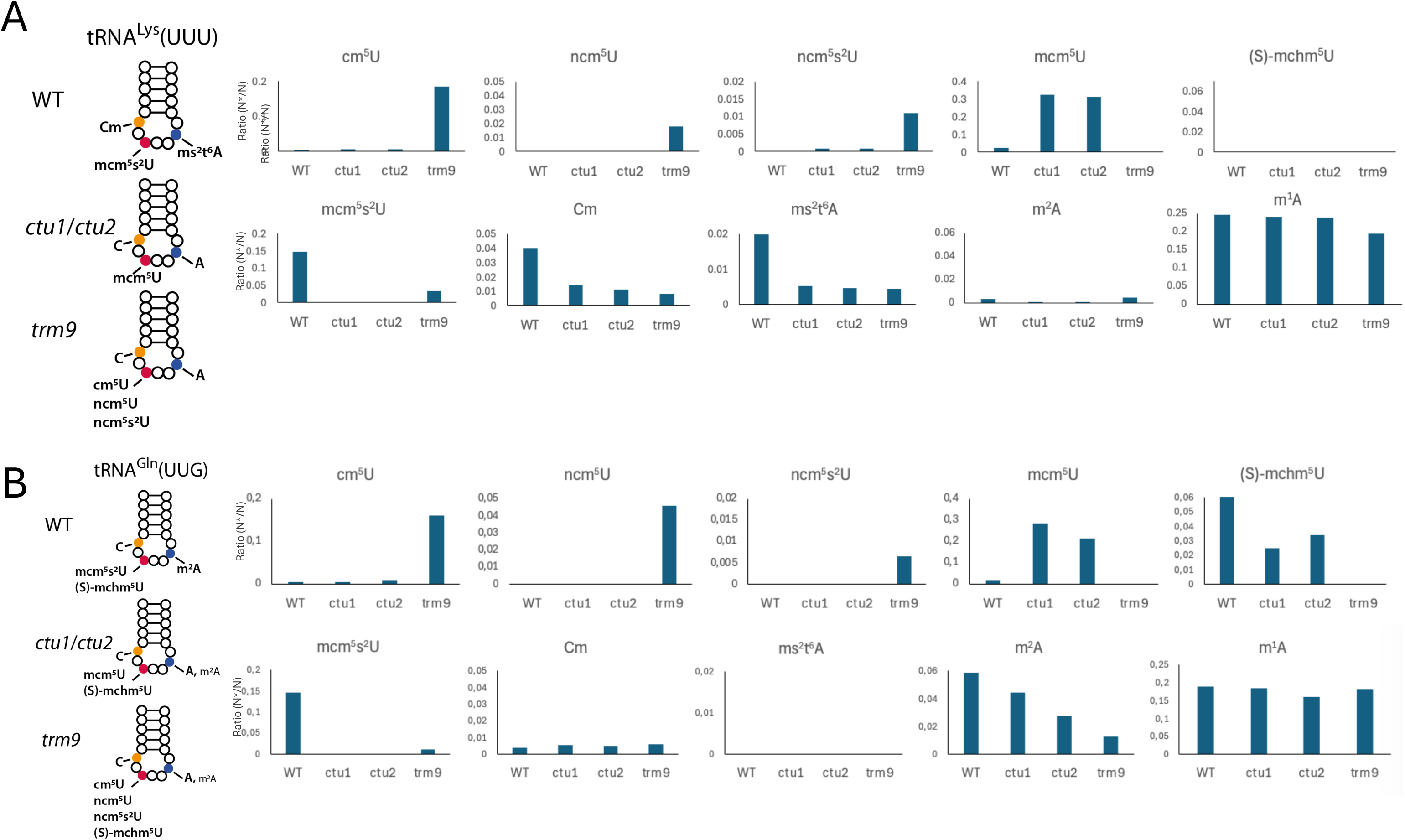
Chemical modifications at sites 32, 34 and 37 for purified tRNA^Lys^(UUU) and tRNA^Gln^(UUG) between Col0 (WT), *ctu1*, *ctu2* and *trm9* backgrounds. Comparison of chemical marks at sites 32 (orange), 34 (red) and 37 (blue) between purified tRNA^Lys^(UUU) (A) and tRNA^Gln^(UUG) (B), in Col0 (WT), *ctu1*, *ctu2* and *trm9* backgrounds. The level of m1A (position 58) is also shown as an example of a modification present in both tRNAs and unaffected in the different backgrounds. Other modifications present in these tRNAs are presented in Supplementary Figure S1. The y-axis of the histogram represents the ratio of the area under the curve (AUC) measured by liquid chromatography/mass spectrometry (LC-MS/MS) of the modified nucleoside (N*) relative to the corresponding unmodified nucleoside (N).

Surprisingly, our mass spectrometry analysis suggests that these two tRNAs known to be substrates of the CTU1/CTU2 complex nevertheless possess different chemical decorations in their anticodon loop. In a first attempt to confirm this result, we used a mim-tRNAseq approach^48^, which can reveal the presence of tRNA chemical modifications, when they interfere with the reverse transcription step. For tRNA^Lys^(UUU), we observed insertions/deletions corresponding to position 32 and misincorporations corresponding to position 37 in the WT, but not in the *ctu1* or *ctu2* mutant background (Figure 3A and Figure S2, note that the modification in position 34 cannot be detected by the mim-tRNAseq). No such errors were detected in tRNA^Gln^(UUG) and tRNA^Glu^(UUC) (Figure S2). These results support our previous conclusion that tRNA^Lys^(UUU) has Cm at position 32 and ms^2^t^6^A at position 37, and that both modifications are almost completely lost in the *ctu1* or *ctu2* thiolation mutants. It also confirms that tRNA^Glu^(UUC) and tRNA^Gln^(UUG) do not have Cm at position 32 and ms^2^t^6^A at position 37 (m^2^A cannot be detected by this approach because it does not cause errors during the reverse transcription step). To further confirm the presence of Cm at position 32 of tRNA^Lys^(UUU), we used a RiboMetSeq approach^49^ (Figure 3B). Using this approach, we confirmed that cytosine 32 of tRNA^Lys^(UUU) is 2’-O-methylated.

**Figure 3.**
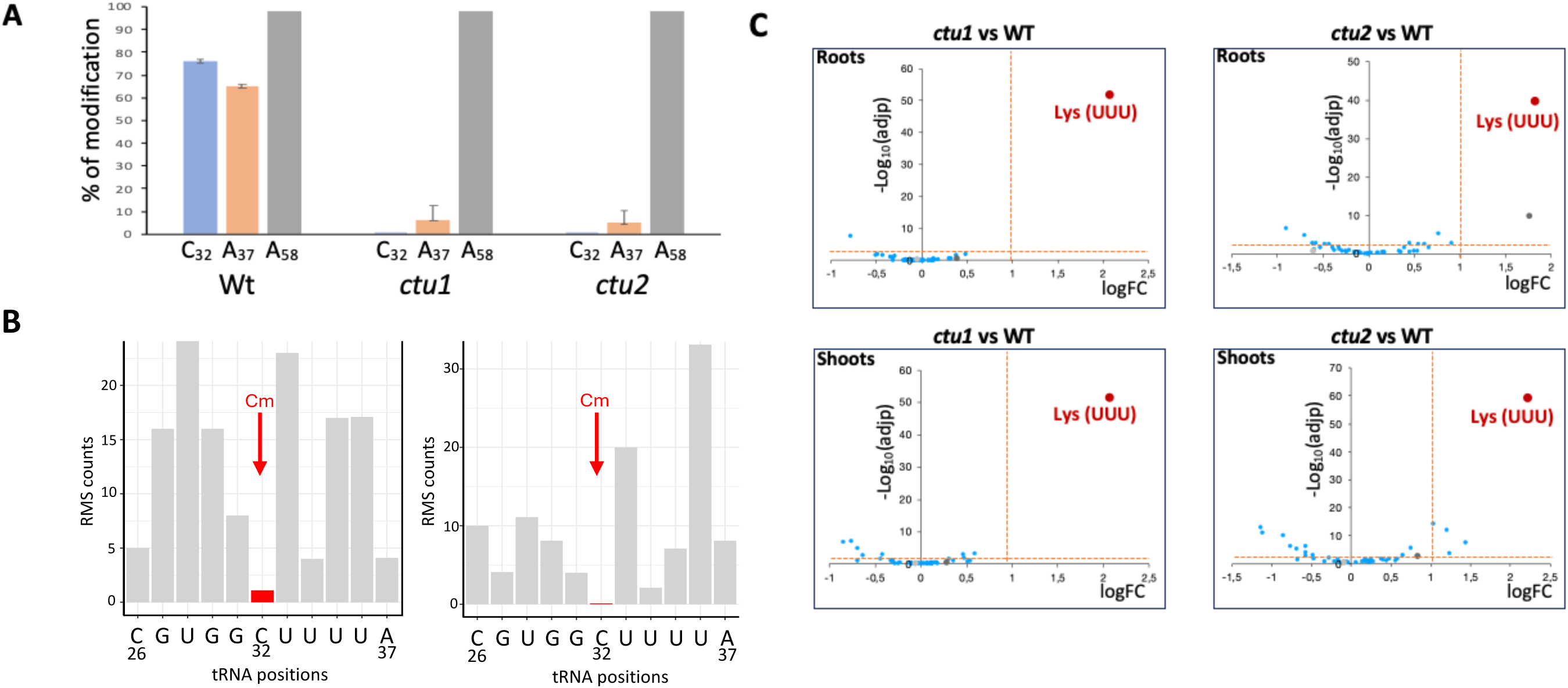
Mim-tRNAseq analysis performed on Col0 (WT), *ctu1* and *ctu2* seedlings. A) Histogram showing the frequency of reverse transcriptase errors at position 32, 37 and 58 of tRNA^Lys^(UUU) in WT (Col0), *ctu1* and *ctu2* mutants. Histogram bars represent the mean of the percentage of errors (n=3) according to mim-tRNAseq analysis. Error bars represent the standard deviation of three biological replicates. The modification at position 32 is detected as insertion/deletion in mim-tRNAseq, while the modification at position 37 and 58 results in missense incorporation. B) RiboMethSeq (RMS) analysis confirming the presence of Cm at position 32 in tRNA^Lys^(UUU). Fragmentation profiles (two independent replicates) from positions 26 to 37 of WT Arabidopsis tRNA^Lys^(UUU) are shown. Red arrows indicate Cm at position 32. C) Volcano plots showing the enrichment in tRNA^Lys^(UUU) in shoots and roots of *ctu1* and *ctu2* mutants compared to WT. The tRNAs from WT and mutant fractions were sequenced and analyzed with mim-tRNAseq, and log2 Fold Change (logFC) were obtained with DESeq2. The statistical confidence of tRNA enrichment is shown on the Y axis as a Log of the adjusted P value. The vertical red dotted lines indicate the threshold for a fold change of 2. The horizontal dotted lines indicate the threshold for an adjusted P value of 0.05. tRNA^Lys^(UUU) is indicated by a red dot.

The genome-wide mim-tRNAseq strategy was also used to document the effect of the *ctu1* and *ctu2* mutations on global steady-state tRNA levels in 15-day-old Arabidopsis shoot and root seedlings (Figure 3C). We observed that steady-state tRNA levels were largely unaffected in the *ctu1* or *ctu2* background, with the clear exception of tRNAs^Lys^(UUU), whose levels are significantly increased (3.5- to 4.5-fold) in both shoots and roots (Figure 3C). This surprising result suggests that the loss of the normal chemical decoration at positions 32, 34, and 37 of the anticodon loop of tRNA^Lys^(UUU) does not destabilize it, but on the contrary likely induces a feedback loop that stimulates the production of new tRNAs to compensate for these epitranscriptomic defects.

### Impact of the *ctu2* mutation on global translation and ribosomes dynamic

A deficiency in one component of the nematode or yeast CTU1/CTU2 complex completely disrupts its thiolase activity^24^. In our case, eliminating the production of either Arabidopsis CTU1 or CTU2 had an almost identical impact on tRNA modifications and steady state levels (see Figures 1 to 3 and S1, S2). Based on these results, we decided to use only the *ctu2* mutant background to assess the impact of tRNA thiolation loss on global translation. To do so, we compared the polysome profiles of WT (Col0) and *ctu2* seedlings (Supplementary Figure S3). Although WT and *ctu2* polysome profiles from 15-day-old roots were similar, the *ctu2* polysome profile from 15-day-old shoots or 5-day-old whole seedlings showed an increase in high-density polysomes compared to WT (Supplementary Figure S3). This difference in profile could result from either an increase in global translation initiation or a decrease in translation elongation rate in the mutant. To distinguish between these two possibilities, we performed a degradome 5’P-seq analysis comparing WT to *ctu2*^50–56^ and used these data to evaluate the capacity of ribosomes to read the AAA, GAA and CAA codons. To do this, we measure the amounts of ribosome-protected fragments in front of all AAA, GAA and CAA codons and compare it to the same situation for their synonymous codons (AAG, GAG and CAG) (Figure 4). The rationale is that if ribosomes slow down when reading AAA, GAA and CAA codons (due to the loss of the mcm^5^s^2^U_34_ modifications (as well as other anticodon loop chemical tags)), we should be able to observe an increase in the protected fragment 16-17 nucleotides in front of these codons (the protected area of a ribosome), as the exoribonuclease will be blocked by ribosomes pausing^50–56^. This effect should not be visible for the synonymous codons because they rely on other isoacceptor tRNAs (with CUU, CUC and CUG anticodon) that are not affected by CTU2 activity. In the meta-analysis of Figure 4A and 4B, we analyzed the accumulation of protected fragment in front of AAA and AAG codons in a WT (Col0) (in red) vs. *ctu2* (in blue) background in shoots and roots. As expected, we observed in all cases that the accumulation of protected fragment follows a three-nucleotide periodicity, which is characteristic of elongating ribosomes transiently protecting translating mRNAs from 5’-3’ exonucleolytic cotranslational degradation^50–56^. A strong increase of protected fragments 16-17 nucleotides before AAA codons can be observed in the *ctu2* background in both shoots and roots. This increase is not present for AAG in the WT and *ctu2* mutant background (Figure 4A). This result shows that the ability of the ribosome to read AAA codons is impaired in *ctu2*, leading to a slowing of translational elongation at these sites and a corresponding accumulation of degradation products. It is interesting to note that this slowing of translational elongation at AAA codons occurs despite the observed 3.5- to 4.5-fold increase in the global amount of tRNA^Lys^(UUU) in the *ctu2* background (see Figure 3C). The same analysis was performed for GAA/GAG (Figure 4C and 4D) and CAA/CAG (Figure 4E and 4F) with similar results but with different intensity. Indeed, the negative effect in the *ctu2* background is stronger for AAA than for GAA, followed by CAA. The global intensity of the minus 16-17 protected fragments is also lower in roots compared to shoots, but always higher compared to the synonymous codons. Based on our polysome profile comparisons and 5’P-seq meta-analyses, we propose that the main effect of the *ctu2* mutation is to slightly impair translational elongation, possibly by slowing down ribosome decoding properties at codons requiring intervention of tRNA^Lys^(UUU), tRNA^Glu^(UUC) and tRNA^Gln^(UUG).

**Figure 4.**
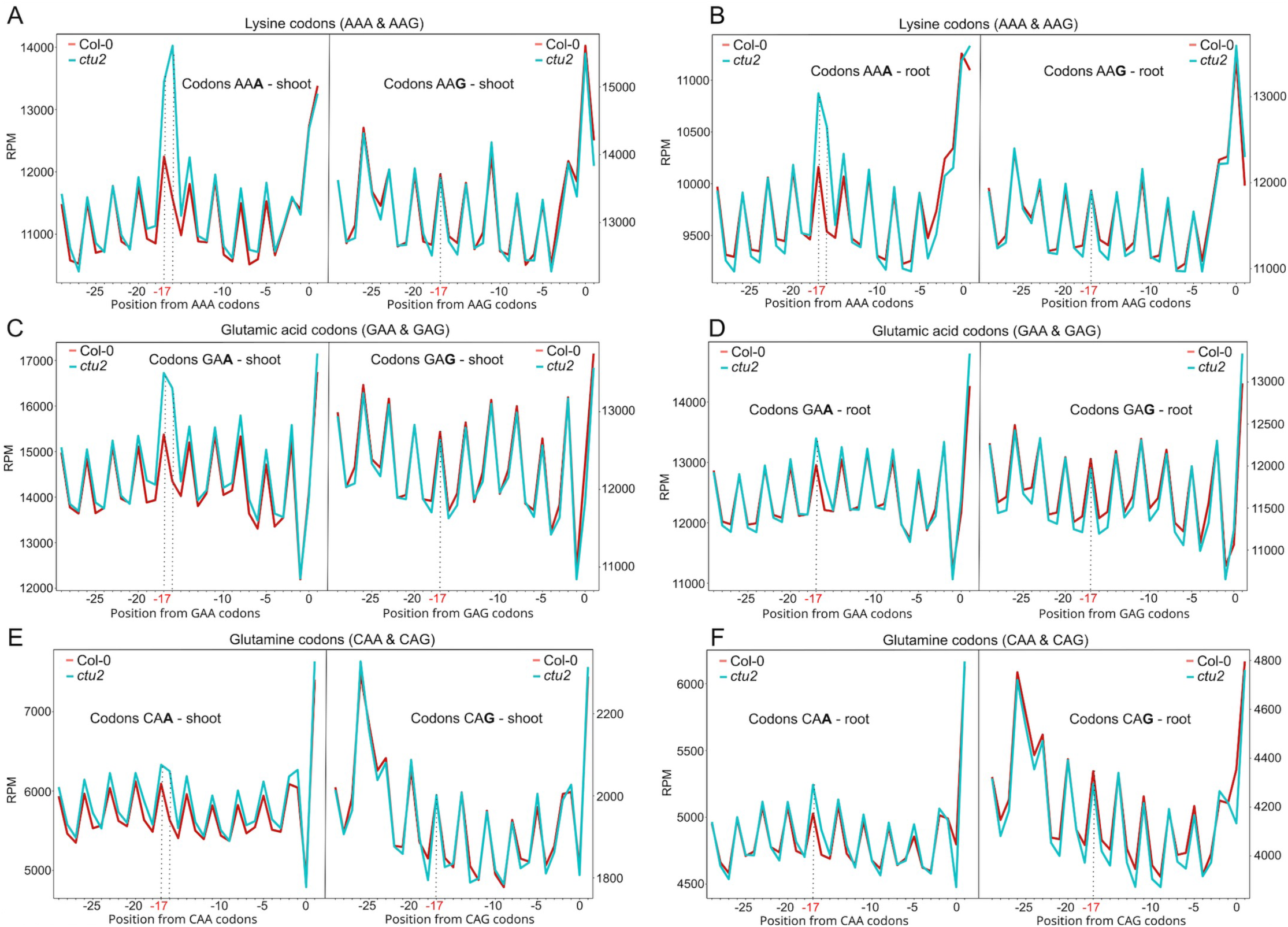
Capacity of ribosomes to read the AAA, GAA and CAA and corresponding synonymous codons in the shoots and roots of WT (Col0) and *ctu2* seedlings. Number of ribosome-protected fragments in front of the AAA and AAG codons in shoots (A) and roots (B). The peak observed at −16 and −17 nucleotides (nt) corresponds to the footprint of the ribosome during the decoding of the related codon^50–56^. For the AAA codon, a pronounced increase of this peak is observed in *ctu2* shoots and roots (light blue) compared to WT (Col0) (red), whereas the synonymous AAG codon does not show this increase (A-B). A similar increase in the intensity of the −16 and −17nt peaks is also observed in front of the GAA and CAA codons in *ctu2* shoots and roots, but not in the corresponding synonymous codons, although with a lower intensity compared to AAA (C to E). All data presented are means of three biological triplicates. RPM = reads per million.

### Impact of the *ctu2* mutation on protein production

We reason that this small but global reduction in translation elongation rate in the mutant should, at least in some cases, result in a corresponding reduction in protein production. Furthermore, we expected that the effect would be stronger for mRNA enriched in AAA, GAA and CAA codons and less important for mRNA depleted in these codons. To test this hypothesis, we performed a quantitative proteomic analysis on 15-day-old Arabidopsis shoot and root seedlings (6 biological replicates) comparing WT (Col0) to the *ctu2* mutant (Figure 5). We observed that 500 and 430 proteins were significantly (padj<0.05) underrepresented in *ctu2* shoots and roots, respectively, compared to WT (Figure 5A). Of these, 170 proteins were underrepresented in both organs. We also observed that 474 and 371 proteins were significantly (padj<0.05) overrepresented in *ctu2* shoots and roots, respectively, compared to WT (Figure 5B). Of these, 172 proteins were overrepresented in both organs. A list of proteins whose steady-state levels are significantly affected in both *ctu2* shoots and roots is presented in Table S3.

**Figure 5.**
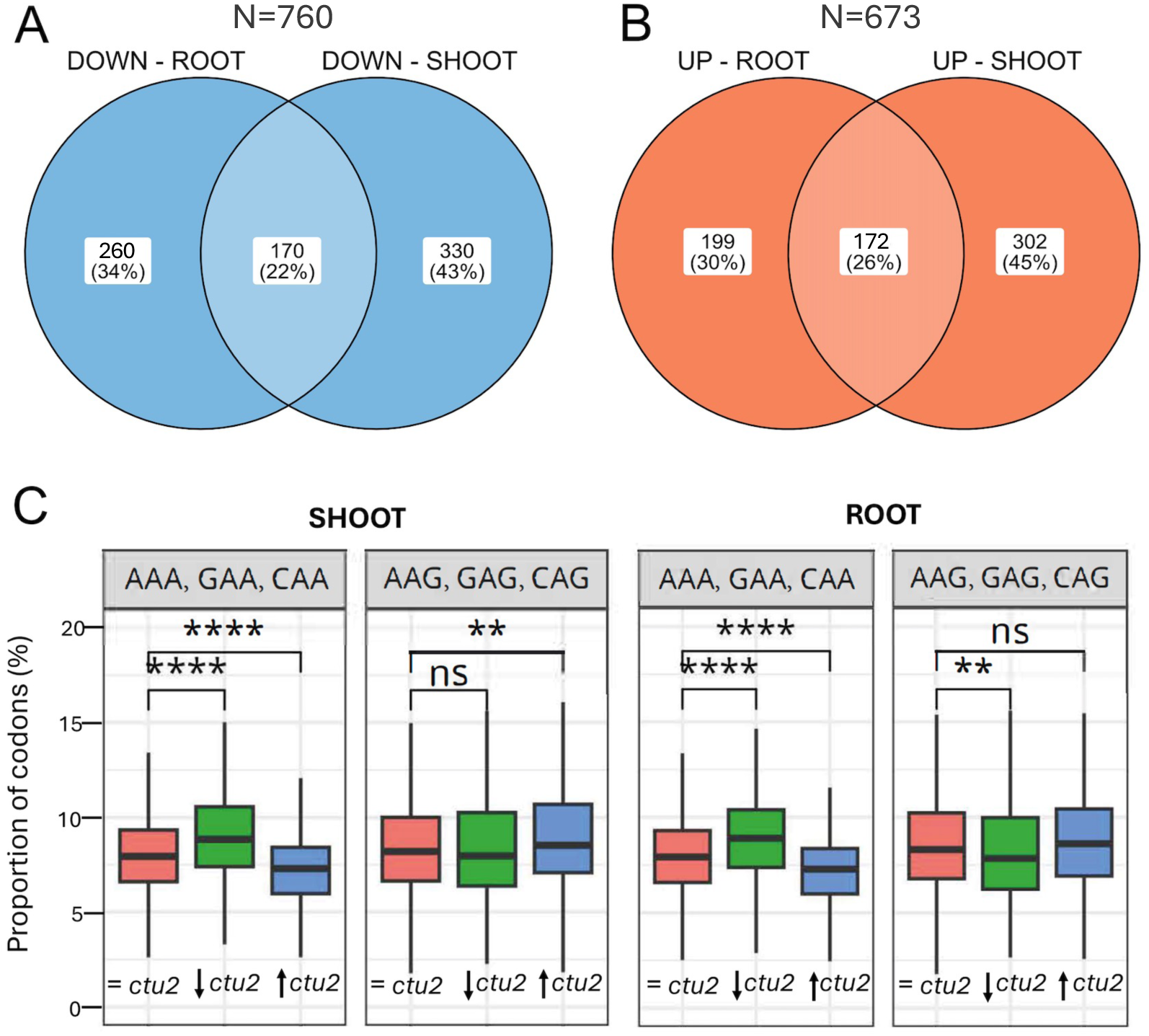
Comparative quantitative proteomic analysis between WT (Col0) and *ctu2* 15-day-old seedlings, shoots and roots. A) Venn diagram showing the number of proteins that are significantly (padj < 0.05) underaccumulated in the shoots and/or roots of *ctu2* compared to the WT. B) Venn diagram showing the number of proteins that are significantly (padj < 0.05) overaccumulated in the shoots and/or roots of *ctu2* compared to the WT. A list of proteins whose steady-state levels are significantly affected in both *ctu2* shoot and root is presented in Table S3. C) Distribution of the percentage of AAA, GAA and CAA or their corresponding synonymous codons AAG, GAG and CAG in nuclear genes encoding proteins with steady-state levels either not affected (=*ctu2*, shoot n=3567, root n=3800), decreased (↓*ctu2*, shoot n=500, root n=430) or increased (↑*ctu2*, shoot n=474, root n=371) in *ctu2* compared to WT in shoot and root (**P<0.01, ****P<0.0001). For each boxplot, the center line represents the median; the lower and upper limit of the boxes represent, respectively, the first and third quartiles.

We next analyzed the proportion of AAA, GAA and CAA or synonymous AAG, GAG and CAG codons in mRNAs encoding proteins that are either down, up or unchanged in the *ctu2* mutant (Figure 5C). We observed that mRNAs encoding proteins that are underrepresented in *ctu2* shoot and/or root are significantly enriched in AAA, GAA and CAA (p<0.0001, non-parametric Wilcoxon test) compared to mRNAs encoding proteins whose level is unchanged. On the contrary, mRNAs encoding proteins overrepresented in *ctu2* shoots are significantly (p<0.0001) depleted in AAA, GAA and CAA, but significantly (p<0.01) enriched in synonymous AAG, GAG and CAG codons compared to mRNAs encoding proteins whose level is unchanged (Figure 5C). The situation is similar in roots, where mRNAs encoding proteins overrepresented in *ctu2* are significantly (p<0.0001) depleted in AAA, GAA and CAA, although they are not enriched in the synonymous codon. This result supports our hypothesis that lower translational elongation efficiency at AAA, GAA and CAA leads to a reduction in protein production mainly for mRNAs enriched in these codons.

### Impact of the *ctu2* mutation on sulfur-related metabolic processes

Next, we performed a Gene Ontology analysis on all proteins that were misregulated in the *ctu2* background compared to WT in shoot (n=974) and root (n=801) and in both organs (n=361) (Figure 6A). In all these cases, the GO term "glucosinolate biosynthesis" was the most highly enriched (12- to 17-fold, Figure 6A), suggesting that the loss of thiolation in three tRNAs affects glucosinolate metabolism. Figure 6B’s heat map shows the shoot and root proteins involved in aliphatic glucosinolate biosynthesis that exhibited significant variations in steady-state levels (*ctu2* versus WT) in our proteomic analysis. We observed a contrasting effect of the *ctu2* mutations on the steady-state levels of these enzymes, as they tend to be overrepresented in the shoot (red) but underrepresented in the roots (blue) (Figure 6B and Table S3). To determine whether these variations in steady-state enzyme levels translate into corresponding changes in glucosinolate concentration, we measured the concentration of glucosinolates in *ctu2* shoots and roots (Figure 7A, B). Indeed, a small increase in the sum of aliphatic glucosinolates was observed in *ctu2* shoots, a result consistent with the observed increase in the amount of the corresponding biosynthesis enzymes. This translated to a greater total glucosinolates content, as the indolic glucosinolates were not affected in the mutants (Figure 7A). However, no variation in glucosinolates levels could be detected in *ctu2* roots, despite a lower amount of the corresponding biosynthesis enzymes (Figure 7B).

**Figure 6.**
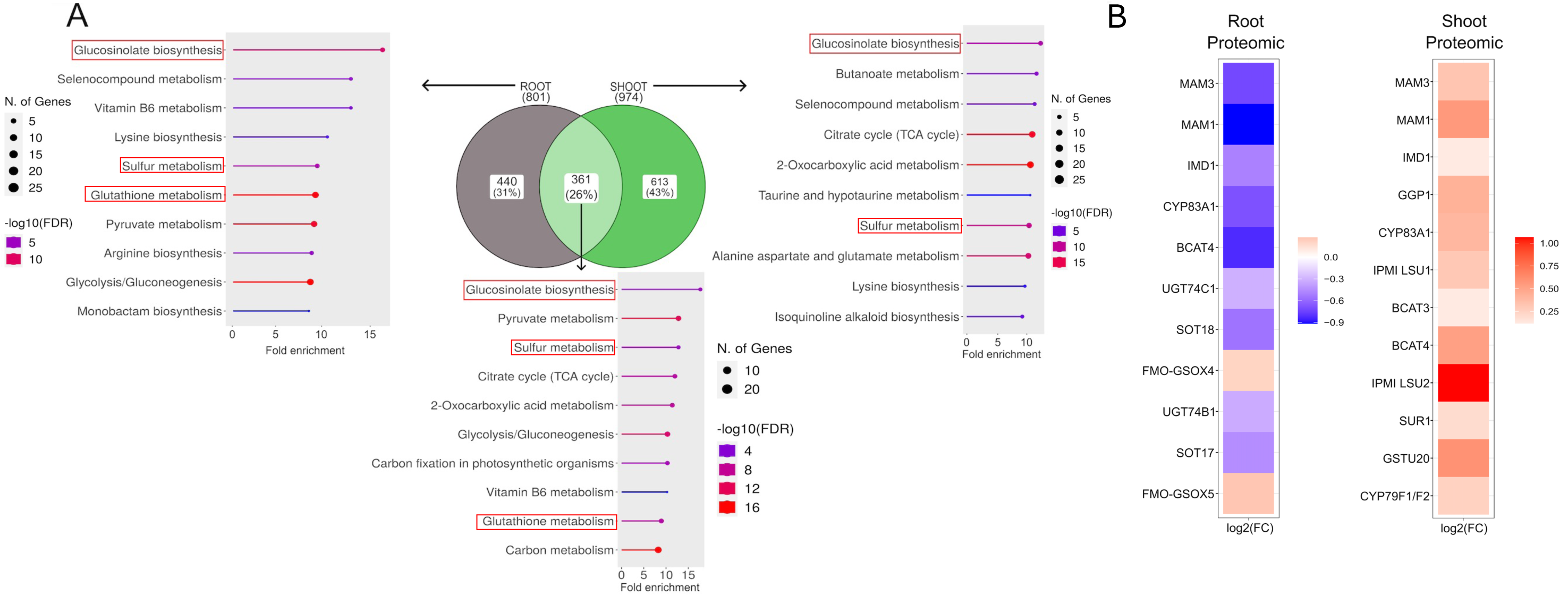
Gene Ontology (GO) analysis of genes encoding proteins with steady-state levels significantly affected in *ctu2* compared to the WT (Col0) in shoot and/or root. A) Most enriched GO terms for proteins misregulated in roots (n=801), in shoots (n=974), or in both tissues (n=361). GO terms in direct relation to sulfur metabolism processes are in red frames. The fold change enrichment is given on the x-axis. The diameter of the final dots is proportional to the number of genes in each category and the significance of each GO-term (-log10(FDR) is color-coded according to the associated legends. B) Heatmap showing the fold change (FC) variation (log2FC) in our proteomic analysis of key proteins involved in aliphatic glucosinolate biosynthesis in *ctu2* shoot or root compared to WT. Negative FC values are shown in blue and positive values are shown in red. See Table S3 for exact FC values.

**Figure 7.**
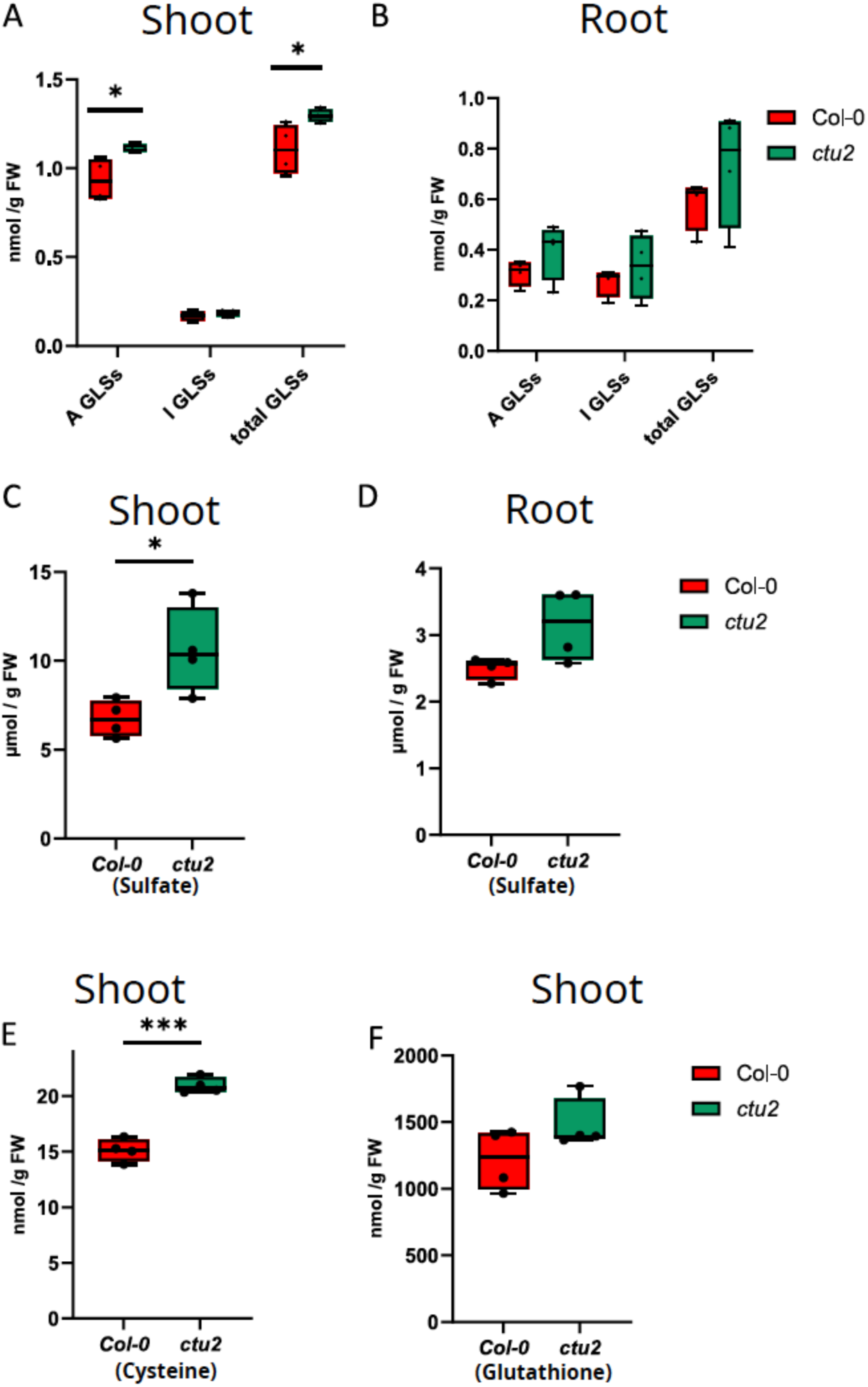
Sulfur-containing metabolites in *ctu2* mutant. Arabidopsis *ctu2* mutants and the WT were grown for 17 days on agarose plates with modified Long Ashton nutrient solution. Aliphatic glucosinolates (A GLSs), indolic (I GLSs), and total glucosinolates (total GLSs) were determined in the leaves (A) and roots (B) by HPLC. Sulfate was measured in the leaves (C) and roots (D) by ion chromatography. Cysteine (E) and glutathione (F) were measured in leaves by HPLC. Asterisks mark values significantly different at *P<0.05 and ***P<0.0001 (T-test, n=4). For each boxplot, the center line represents the mean; the lower and upper limit of the boxes represent, respectively, the first and third quartiles.

In addition, the GO term “glutathione metabolism” and the more general term "sulfur metabolism", were also enriched (more than 10-fold) in proteins misregulated in the *ctu2* background (Figure 6A). This led us to monitor the concentration of sulfate, cysteine (the sulfur donor for tRNAs) and glutathione in the *ctu2* mutant (Figure 7). We observed that both sulfate and cysteine were significantly (p<0.05 and p<0.0001, respectively, T-test n=4) more abundant in *ctu2* shoots, while glutathione was not affected (Figure 7C, E, F). In accordance with no changes in glucosinolates levels in the roots, sulfate levels were also not affected in this organ (Figure 7D). Taken together, these results show that the loss of U_34_ thiolation on tRNAs feedbacks on key sulfur-related metabolic processes, suggesting that the level of tRNA thiolation might function as a sensor to regulate these processes.

## Discussion

In this study, we examined the nature of the chemical marks present at the U_34_ position of Arabidopsis tRNAs. Using total tRNAs, we first demonstrated that three primary modifications occupy this anticodon loop position in wild-type plants: ncm^5^U, the most abundant, followed by mcm^5^s^2^U and (S)-mchm^5^U (Figure 1). It is widely accepted that a given tRNA species possesses only one of these U_34_ modifications, and that different U_34_ modifications are found exclusively on different tRNA species^26^. For example, although ncm^5^U is the most common U_34_ modification overall (Figure 1), it is absent from tRNA^Lys^(UUU) and tRNA^Gln^(UUG) (Figure 2) and therefore must decorate other Arabidopsis tRNAs with a U in position 34. However, we provide here evidence that Arabidopsis tRNA^Gln^(UUG) can exist in wild type plants with either mcm^5^s^2^U_34_ or (S)- mchm^5^U_34_ (Figure 2B). We further observed that the thiolation mark of tRNA^Gln^(UUG) is completely lost in the *ctu2* mutant, though part of the (S)-mchm^5^U modification is retained (Figure 2B). In addition, we saw that the *ctu2* mutation has a much smaller impact on CAA decoding compared to AAA decoding, which uses tRNA^Lys^(UUU) that do not have (S)-mchm^5^U in this background (Figures 2A and 5). Based on these observations, an intriguing possibility is that tRNA^Gln^(UUG) decorated with (S)- mchm^5^U_34_, instead of mcm^5^s^2^U_34_, could also facilitate CAA decoding in wild type plants. Alternatively, these differences in decoding properties could result from the different modifications found in position 37 for these two tRNAs: m^2^A for tRNA^Gln^(UUG) and ms^2^t^6^A for tRNA^Lys^(UUU), as they could influence differently CAA and AAA decoding^26,57^. In any case, the possibility that a given tRNA can have different U_34_ chemical modifications is an unexpected new layer of epitranscriptomic complexity that will need to be further evaluated.

Until recently, only a limited number of cases of tRNA modification interdependency have been described. Examples include several crosstalk occurring during the maturation of yeast tRNA^Phe^ ^58^, between queuosine (Q_34_) and m^5^C_38_ in several *S. pombe* and human tRNAs^59,60^, and m^3^C_32_ and i^6^A_37_ in a subset of *S. pombe* tRNAs^Ser^ ^61^. The introduction of the SingLe-read Analysis of Crosstalks (SLAC) method recently allowed for the detection of large number of new crosstalk situations in yeast, mouse, and human tRNAs^62^. For instance, new crosstalks were identified between m^1^G_9_ and m^1^A_58_ in yeast tRNA^Lys^(CUU), as well as between m^2^_2_G_26_, m^1^I_37_, and m^1^A_58_ in the human isodecoders tRNA^Ala^(AGC)-4/8, among many others^62^. The SLAC approach revealed that the nature and intensity of crosstalk vary among tissues and under stressful conditions. Additionally, it showed that crosstalk influences the binding of amino acids to tRNAs and translation efficiency. These results suggest that tRNA modification interdependencies represent a new layer of complexity with physiological importance that affects the tRNAome and, ultimately, translation^62^. In this study, we describe an unreported tRNA crosstalk involving Cm_32_, mcm^5^s^2^U_34_, and ms^2^t^6^A_37_ in the Arabidopsis tRNA^Lys^(UUU). This crosstalk is not detectable using the SLAC method because SLAC only identifies modifications that generate a mismatch during reverse transcription, and this is not the case for Cm or mcm^5^s^2^U^62^. The highly conserved nature of the mcm^5^s^2^U_34_ modification suggests that this network may exist in other eukaryote species as well but has gone unnoticed thus far due to the technical limitations of the SLAC approach.

The loss of all three chemical modifications at positions 32, 34, and 37 of tRNA^Lys^(UUU) in *ctu1* or *ctu2* backgrounds strongly influenced the steady-state level of this tRNA. Indeed, we observed a 3.5- to 4.5-fold increase in the amount of tRNA^Lys^(UUU) in both shoots and roots (Figure 3C). This is unusual because the loss of tRNA chemical decorations typically either results in the destabilization of the corresponding tRNAs and a reduction in their steady-state level or have no effect on stability and tRNA level^63^. It seems unlikely that the increase in tRNA^Lys^(UUU) is due to an improved maturation process and/or stabilization of the mature, yet chemically deficient, tRNAs. It is tempting to speculate, however, that the plant stimulates the production of this tRNA family to compensate for the decoding defect due to the loss of the three chemical marks. *Arabidopsis thaliana* produces its cytoplasmic tRNA^Lys^(UUU) from thirteen interspersed nuclear genomic loci, three of which can be distinguished as having sequence variations (http://plantrna.ibmp.cnrs.fr/). Our mimtRNA-seq data represent this genomic diversity, indicating that at least four of the thirteen loci might be targeted by this putative transcriptional activation. A previous study found no methylation mark at any of the thirteen tRNA^Lys^(TTT) loci^64^. Therefore, if this feedback is based on an epigenetic mechanism, it must act directly at the chromatin level. An example of such epigenetic regulation of tDNA genes was observed in *Bombyx mori*, where the positioning of a nucleosome near an upstream *cis*-element was found to stimulate transcription of a glycine tDNA gene^65^. Alternatively, transcription could be stimulated by the production of a *trans*-acting regulator. However, this putative regulator would need to act preferentially on tRNA^Lys^(TTT) loci since the loss of thiolation does not significantly impact other tRNA gene families. It would also need to target multiple members of this gene family. Further work is needed to elucidate this intriguing observation.

As expected, the loss of U_34_ thiolation for three tRNA species, has almost no impact on global translation. We were nevertheless able to document a small increase in high polysomes content in 5-day seedlings and 15-day shoots (Supplementary Figure S3), likely due to a slowdown of translation elongation at AAA, GAA and CAA codons (Figure 4). Recently, the ribosome profiling technique also revealed that yeast ribosomes have difficulty decoding these three codons following the loss of U_34_ thiolation, though the intensity of the impact (AAA > CAA > GAA) differs from that reported in here in Arabidopsis (AAA > GAA > CAA)^66^. In addition, we observed that the difficulty in reading these three codons limits the production of protein coded by mRNAs that are enriched in these codons more severely (Figure 5). Finally, we observed that loss of tRNA thiolation directly or indirectly increases or decreases the accumulation of many proteins involved in several major sulphur-related metabolic processes. (Figure 6 and 7). Overall, these results support the hypothesis that tRNA modification dynamics are a general mechanism by which metabolic production, tRNA function, translation, and protein synthesis are adjusted to maintain cell homeostasis under normal growth conditions^19,27^. Thus, modification of tRNA epitranscriptomic marks might represent a mechanism by which the global nutrient response and perception of metabolic signals are integrated with translation. This hypothesis, essentially developed on the basis of evidence obtained in unicellular organisms (yeast and bacteria), is supported by the need for key metabolites to modify tRNA (such as cysteine, SAM, and acetyl-CoA to generate mcm^5^s^2^U and threonine to generate ms^2^t^6^A) and the observation that differently modified tRNA can direct translation towards different mRNAs depending on their codon usage. We provide here further support for this hypothesis, this time using a multicellular organism. We show that preventing the thiolation of three tRNAs feeds back on sulfur-compound metabolism, leading to differential accumulation of many enzymes involved in these processes (Figure 6 and Table S3) which further affects concentrations of key sulfur-containing metabolites (glucosinolates, sulfate, and cysteine) (Figure 7). The moderate increase in the concentration of aliphatic glucosinolates in *ctu2* shoot and its stability in *ctu2* roots is intriguing as it does not fully correlate with variation in protein levels (1.5 to 2 folds, up in *ctu2* shoot and down in *ctu2* roots). One possibility is that glucosinolates are reallocated from shoots to roots in the mutants as long-distance transport of these molecules was shown to occur in Arabidopsis^67^. Another possibility would be an adjustment of catabolic processes to maintain glucosinolates concentrations despite variations in biosynthesis. Interestingly, proteins involved in the TCA cycle, pyruvate metabolism, glycolysis, and lysine biosynthesis are also significantly overrepresented in our proteomic data (Figure 6A and Table S3) and identified as proteins misregulated in the *ctu2* mutant (see Table S3). This suggests that variations in tRNA thiolation level could more generally affect the global nutritional status of the plant and help to adapt the metabolic rate to protein synthesis.

## Methods

### Plant material and growth conditions

For tRNA extractions, polysome profiling and NGS analysis, plants were grown *in vitro* for 5 days or 15 days on synthetic Murashige and Skoog (MS) medium (MS0213, Duchefa) containing 1% sucrose and 0.8% plant agar at 20°C under a 16h light (120 µmol.m-2.s-1)/8h dark cycle. The wild-type Columbia (Col0) ecotype was used with the following Col0 tDNA insertion transgenic lines obtained from the Eurasian Arabidopsis Stock Centre (NASC): *drl1*: SALK_140551C, *trm9*: SALK_135308C, *ctu1*: GK-709D04, *ctu2*: GK-686B10, *alkbh8*: SALK_094502C. For sulfur-containing metabolites analyses, Arabidopsis *ctu2* mutants and the WT were grown for 17 days on agarose plates with modified Long Ashton nutrient solution.

### Total RNA extraction and tRNA purification

Total RNA was extracted from 50 mg of Arabidopsis tissue using the Monarch Total RNA Miniprep Kit (NEB, T2010S). The quantity and quality of the RNA were assessed using the Qubit and the Agilent 2100 Bioanalyzer with a Plant RNA Nano Chip, respectively (Invitrogen). Only total RNA with an RIN greater than 8 was used for further experiments. To purify the total tRNA, 8 µg of total RNA was mixed with one volume of 2X RNA loading buffer (95% formamide, 18 mM EDTA, 0.025% SDS, and 0.025% of xylene cyanol) and loaded onto a 15% acrylamide gel containing 8 M urea and 1X TBE buffer. The gel was preheated at 65 °C and pre-run for 15 minutes at 250 V. The total RNA was denatured for two minutes at 70°C and then directly loaded onto the polyacrylamide gel (without a previous incubation on ice). The gel was run for 3.5 hours at 100 V to separate the RNAs. For staining, the gel was placed in a 50 mL, 0.5X TBE bath containing 5 µL of SybrGold (10,000X concentrated, ThermoFisher Scientific) and shaken slowly for 10 minutes. The gel was washed in a 50 mL, 0.5X TBE bath. The tRNAs were visualized under UV light and excised from the gel at the corresponding size (65–94 nucleotides) using a sterile scalpel. The tRNAs were recovered using the Zymo Research ZR Small RNA PAGE Recovery Kit (R1070) and eluted in 11 µL of RNase-free water. The tRNA concentration was measured using the Invitrogen RNA High Sensitivity Qubit Assay. The tRNAs were stored at −80 °C prior to LC-MS/MS analysis.

For tRNA^Lys^ and tRNA^Gln^ purification, total RNA was extracted from 15-day-old in vitro-grown plants using Tri-Reagent (Sigma-Aldrich) and enriched in small RNAs (<150 nt) by LiCl precipitation, as described previously^68^. Briefly, 100 µg of small RNA were mixed with 10 µL of specific biotinylated oligonucleotide (100 µM) in 500 µL of 5X SSC, 0.05% (w/v) SDS. After denaturation (5 min at 70 °C) and rapid cooling (30 s on ice), hybridization was performed for 3 h at 45 °C. Streptavidin Sepharose High Performance beads (100 µL, GE Healthcare) were loaded onto 35 µm receiver columns (Macherey-Nagel) and washed twice with 500 µL of 5x SSC, 0.05% SDS, with centrifugation (15 s, 11,000g) between washes. The hybridization mixture was loaded onto the column and incubated for 15 min at room temperature followed by 15 min at 45 °C with gentle stirring. After centrifugation (15 s, 11,000g) and discarding of the flowthrough, the column was washed sequentially with: 500 µL 2x SSC for 5 min at 45 °C, 500 µL 2x SSC for 15 min at 48 °C, and 500 µL 2x SSC for 15 min at 52 °C. Bound tRNAs were eluted with 400 µL TE-urea (5 min, 60 °C) followed by 200 µL TE-urea (5 min, 60 °C). Eluates were pooled, ethanol-precipitated overnight at –20 °C in the presence of 0.5 µL glycogen, and centrifuged (30 min, 16,000g). The resulting tRNA pellet was resuspended in 10 µL water and quantified using a Nanodrop spectrophotometer. The sequences of the two biotninylated synthetic oligonucleotides (Eurofins) used for tRNA^Lys^(UUU) and tRNA^Gln^(UUG) purification were the following: TGGCGCCGTCTGTGGGGATCGAACCCACGGCCACGTG/3Bio/3’ TGGAGGTTCTACCGGGAGTCGAACCCAGGTCGC/3Bio/ −3’

### Construction of the *ctu2*:pUBQ-CTU2 complemented line

The chemically synthesized CTU2 coding sequence was cloned into a modified pBluescript entry vector containing a chloramphenicol resistance gene. Then, it was recombined using the Gateway site-specific recombination (SSR) cloning system (Invitrogen, CA) into the binary Gateway overexpression vector pUB-DEST, which contains the ubiquitin promoter. The tDNA insertion line GK-686B10, mutated for CTU2, was transformed using a recombinant *Agrobacterium tumefaciens* culture and the floral dip technique^69^. The expression level of CTU2 in the resulting BASTA-resistant transgenic lines was monitored by quantitative reverse transcription PCR (qRT-PCR).

### Nucleosides detection by LC-MS/MS analysis

For the tRNA digestion, 60 ng of tRNA or 10 ng two different purified tRNAs, tRNAs^Lys^(UUU) and tRNA^Gln^(UUG) were diluted to a final volume of 20 uL using ultra-pure water. tRNA were digested with 1U of nuclease P1 from *Penicillum citrinum* (Sigma-Aldrich, Cat No. N8630-5X1V) in a 100 mM solution of ammonium acetate adjusted to a final pH of 5.3 with glacial acetic acid. Nucleotide digestion was performed by incubating at 42°C for 2h. For nucleotide phosphate removal, 1U of alkaline phosphatase from *Escherichia coli* (Sigma-Aldrich, Cat No. P4252-100UN) was added in a solution of ammonium acetate to a final concentration of 1M at pH 7.0. Each sample was incubated at 37°C for 2h. Finally, prior to LC-MS/MS analysis, 60 uL of mobile phase A (0.5% acetic acid in water ULC/MS) was added to each sample, followed by filtration with a 0.22 µm filter (Merck, Cat No. SLGVR04NL) into LC-MS/MS vials (Agilent Cat No. 5188-2788) and injection of 5 µL into the liquid chromatography/mass spectrometer.

For the LC method, a Shimadzu Nexera LC-40 system was used. Chromatographic separation of nucleosides was performed at 35°C in a C18 column (Synergi Hydro-RP; 4 µm particle size, 250 mm x 2.0 mm, 80 Å, Phenomenex, Cat No: 00G-4375-B0) using the solvents 0.5% acetic acid in water ULC/MS and pure acetonitrile as mobile phase A and B, respectively. The flow rate was set at 0.4 mL/min and gradient elution was performed, starting with 100% phase A for 3 minutes, followed by an increase in phase B up to 8% for 10 minutes and up to 40% for another 10 minutes. After 23 minutes of run time, the column was rinsed with 80% phase B for 7 minutes and re-equilibrated with 100% phase A for an additional 5 minutes to establish initial conditions.

For the LC-MS/MS nucleoside analysis, a triple quadrupole mass spectrometer NX-8060 (Shimadzu) coupled to an electrospray ionization (ESI) source, was used. Instrument settings for the ESI source are described in Table S4A. Forty-four nucleosides were selected and analyzed in positive ion mode using multiple reaction monitoring (MRM) acquisition mode (Table S4B). The MRM transitions and specific MS parameters are described in Table S4B. For each target, the retention time window and dwell time, were set to 3 min and 20 ms, respectively. The total area under the curve (AUC) for each modified and unmodified nucleoside was extracted by LabSolution Insight software version 3.8 SP3 (Shimadzu). For the qualitative quantification, the ratio of the AUC of the modified nucleoside relative to the AUC of the corresponding non-modified nucleoside was calculated. Standard nucleosides were purchased from Biosynth (Bratislava, Slovakia), with the exception of (R)- and (S)-mchm^5^U nucleosides, which were supplied by MedChemTronica (Sweden).

### Mim-tRNAseq, RiboMet seq and degradome (5’P-seq) analysis

For the mim-tRNAseq analysis, biological triplicates of total tRNAs from Arabidopsis WT and T-DNA insertion mutants were extracted as described above for tRNA^Lys^ and tRNA^Gln^ purification. They were sequenced using an adaptation of the mim-tRNAseq protocol^48,70^ as described in^71^. RiboMet-seq analysis was carried out by the Epitranscriptomics and Sequencing (EpiRNA-Seq) Core Facility, University of Lorraine (Nancy, France), as described previously^49^, using the same tRNA samples as for mim-tRNAseq. Degradome (5’P-seq) libraries were prepared as previously described^56^. Briefly, 50 µg of total RNA was subjected to two rounds of poly(A+) purification. After ligation of 5ʹ adaptors, excess adaptors were removed by a new round of poly(A+) purification. Reverse transcription was performed using the SuperScript IV system according to the manufacturer’s instructions. cDNAs were amplified by 11 cycles of PCR. Libraries were purified using SPRIselect beads prior to quality control and normalization. Library quality was checked using an Agilent High Sensitivity DNA Kit (Agilent). Libraries were normalized, multiplexed and sequenced on NextSeq 550 (Illumina) in 75 pb single reads. After sequencing, raw reads for Read 1 were used and trimmed to 50 pb before mapping. Metagene analysis was performed using FIVEPSEQ software with standard parameters^72^. To generate metagene profiles around start, stop and individual codons, “meta_counts_start”, “meta_counts_term” and “codon_paused” generated by FIVEPSEQ were used.

### Polysome gradients

Polysome profiling was performed as described previously^73^. For polysome quantification, 100 mg ground powder (5-day-old seedlings or 15-day-old shoot or root) was incubated for 10 min on ice with 3 volumes of lysis buffer (200 mM Tris-HCl, pH: 9, 200 mM KCl, 25 mM EGTA, 35 mM MgCl2, 1% (v/v) detergent mixture [1% (v/v) Tween 20, 1% (v/v) Triton, 1% (w/v) Brij35 and 1% (v/v) Igepal], 1% (w/v) sodium deoxycholate, 0.5% (w/v) polyoxyethylene tridecyl ether, 5 mM dithiothreitol, 50 μg.mL^-1^ cycloheximide, 50 μg.mL^-1^ chloramphenicol and 1% (v/v) protease inhibitor cocktail (Sigma-Aldrich)). The extract was centrifuged at 16,000g for 10 min, and 300 µL of the supernatant was loaded onto 9 mL 15-60% sucrose gradients. Ultracentrifugation was performed at 38,000 rpm (180,000g) for 3 hours using an SW41 rotor (Beckman Coulter). Polysome profiles were recorded using an ISCO absorbance detector at 254nm, analyzed using the PeakTrack software and plotted using Excel. The total amount of ribosomes was evaluated by measuring the area between the 40S, 60S, monosomes and polysomes profile and the basal lines, while the number of polysomes was evaluated by measuring the area on the low or high polysome portion of the profile only. Values represent the mean of three biological replicates.

### Protein purification and proteomics study

For protein purification, roots and shoots samples from 15-day-old seedlings were harvested and ground separately in liquid nitrogen. Total proteins were extracted using a protocol adapted from Hurkman and Tanaka^74^. Protein pellets were resuspended in 1% SDS. Protein concentration was assessed using a Qubit Protein Assay Kit (Invitrogen). The leaf and root proteomic preparation was carried out using the SP3 automated protocol on the LT Bravo Liquid handler (Agilent technologies). Briefly, 5 µg of total protein from each sample were diluted in Sodium Dodecyl Sulfate (SDS) to a final percentage of 1%, reduced with 5 µL of Dithiothreitol (DTT) at 80 mM for 30 min at 60°C, and then alkylated with 5 µL of Iodoacetamide (IAA) at 200mM for 30 min at 30°C. Protein capture was performed by adding 5 µL of Cytiva Sera-Mag Speedbead at 100 mg/mL (1:1 mix of E3 and E7 magnetic particles) and 35 µL of acetonitrile. Proteins are left to bind to the beads while shaking for nine consecutive repetitions of 30 seconds at 1500 rpm and 90 seconds at 100 rpm. Beads were then washed twice with 80% ethanol and once with 100% acetonitrile. Digestion was performed by adding 250 ng of trypsin/lysC mixture in 40 µL of 50mM ABC and incubating the sealed plate overnight at 37°C and 450 rpm and was stopped with 10 µL of 5% formic acid. Peptide recovery was carried out using a magnetic rack. The peptide concentration was quantified using the NanodropOne Microvolume UV-Vis Spectrophotometer (reference ND-ONE-W, ThermoFisher Scientific). A total of 400 ng of peptides were loaded onto Evotips Pure™ according to manufacturer’s instructions. An EvoSep One (EvoSep) was used as liquid chromatography system with a PepSep C18 column, 15 cm x 150 µm x 1.5 µm (Bruker Daltonics). The elution was carried using a 30 samples per day (SPD) method, which corresponds to a 34-minutes gradient, using 0.1% FA in water as mobile phase A (Biosolve) and 0.1% FA in acetonitrile as mobile phase B (Biosolve). The EvoSep One was coupled to a trapped ion mobility spectrometry quadrupole time-of-flight mass spectrometer (timsTOF HT, Bruker Daltonics). Data from all samples were acquired in a Data-Independent Acquisition (DIA) mode. Data-Dependent Acquisition (DDA) was used on the same samples to generate a library. For both methods, the conditions were the following: capillary voltage 1100 V; dry gas 3.0 L/min and dry temperature 180°C. MS1 and MS2 spectra were collected in the range of 100 to 1700 m/z. The collision energy was set by linear interpolation between 59 eV at an inverse reduced mobility (1/K0) of 1.60 Vs/cm^2^ and 20 eV at 0.6 Vs/cm^2^. For DIA, the accumulation and ramp times were 75 ms. The diaPASEF window scheme was in dimensions from m/z 340 to 1100 and in dimension 1/K0 from 0.68–1.33. For DDA, the accumulation and ramp times were 100 ms. Three ions of the Agilent ESI-Low Tuning Mix were used to calibrate the ion mobility dimension (m/z [Th], 1/K0 [Th]: 622.0289, 0.9848; 922.0097, 1.1895; 1221,9006, 1.3934). DDA-PASEF parameters included the following: mass/charge ratio (*m/z*) range: 100 to 1700, mobility (1/K0) range: 0.75 to 1.25 V⋅s/cm^2^, the accumulation and ramp time were 100 ms. Target intensity per individual PASEF precursor was set to 20000. The values for mobility-dependent collision energy ramping were set to 59 eV at an inversed reduced mobility (1/K0) of 1.60 V·s/cm^2^ and 20 eV at 0.6 V·s/cm^2^. Collision energies were linearly interpolated between these two 1/K0 values. LC-MS acquisitions were performed in combination with the PaSER (Parallel Search Engine in Real-time; Bruker Daltonics) platform that enables real-time processing of PASEF DDA and dia-PASEF spectral data streamed from the timsTOF Pro to the PaSER box. The Uniprot database of *Arabidopsis thaliana* was used as reference (Release_2024_04). The library contains 5712 protein groups, 76264 precursors in 70654 elution groups. The parameters used were the following: trypsin as digestion enzyme, maximum 1 missed cleavage, minimum 7 amino acids and maximum 30 amino acids for peptide size, minimum 1 and maximum 4 for precursor charge range, minimum 300 and maximum 1800 for the precursor m/z. To increase identification confidence, the TIMScore implemented in PaSER was used, which compares experimental collision cross section (CCS) values with predicted CCS values for candidate peptides. Some modifications, induced by the sample preparation protocol, were studied: methionine oxidation as variable modifications and cysteine carbamidomethylation as fixed modification. The maximum number of variable modifications was set to 1. Mass tolerances were set to 20 ppm for precursor ions and 30 ppm for fragment ions. A protein identification FDR was set at 1% and library was generated with tsv export from paser-box. Protein quantification was then performed using the DIA-NN software (version 1.9.1) with a library based on DDA runs. Mass accuracy and MS1 accuracy were determined automatically. Peptidoforms, heuristic protein inference, and QuantUMS (high precision) were used. LFQ intensities obtained from DIA-NN software (version 1.9.1) were processed using the DIA-Analyst platform (version 0.9.2). Proteins considered contaminants and redundant were removed, and only proteins present in 100% of samples were retained. LFQ data for each protein were transformed using the log_2_(x) formula and grouped according to genotype (Col0, *ctu2*). Missing values were not imputed. An adjusted p-value threshold of 0.05 (t-statistic correction) was applied to determine which proteins were significantly regulated in each comparison.

### Codon composition and Gene Ontology analyses

We calculated the percentage of AAA, GAA and CAA codons and their corresponding synonymous codons AAG, GAG and CAG in nuclear Arabidopsis protein encoding genes using the Araport11 database (https://www.araport.org/) and established their distribution in the different categories (proteins not affected in *ctu2*, decreased in *ctu2* and increases in *ctu2*). For each distribution comparison, we performed a non-parametric Wilcoxon test on the codon proportion distributions. The gene ontology analyses were done using the ShinyGo (version 0.82) website^75^.

### Analysis of sulfur-containing metabolites of sulfate, cysteine and glucosinolates

The concentrations of sulfur-containing metabolites were determined as described previously^76^. Briefly, glucosinolates were extracted from ca. 25 mg leaves and roots in hot 70% methanol with 2.5 nmol sinigrin as internal standard. After incubation at 70 °C for 45 min and centrifugation, the extracts were loaded onto DEAE Sephadex A-25 columns and washed twice with water and 0.02 M sodium acetate, before 75 µl sulfatase solution (30 U) was added to the column. After overnight incubation the resulting desulfo-glucosinolates were eluted with sterile water and resolved by HPLC (Spherisorb ODS2, 250 3 4.6 mm, 5 μm; Waters) using a gradient of acetonitrile in water. The desulfo-glucosinolates were detected by UV absorption at 229 nm, and quantified using the internal standard and response factors^76^. Sulfate was extracted from the leaves and roots in water and determined by Dionex ICS-1100 ion chromatography system using a Dionex IonPac AS22 RFIC 4x 250 mm analytic column (Thermo Scientific)^76^. The thiols, cysteine and glutathione, were measured in leaves after extraction in 0.1 M HCl, reduction with DTT, and derivatisation with monobromobimane. The thiol conjugates were separated by HPLC on Spherisorb™ column (ODS2, 250 x 4.6 mm, 5 µm, Waters), using a gradient of methanol in 0.25% acetic acid, and detected fluorimetrically with a 474 detector (excitation, 390 nm, emission, 480 nm)^76^.

## Data Availability

Data supporting Figure 1 and 2 (tRNA mass spectrometry) can be found in Table S1. Mim-tRNAseq datasets supporting Figure 3 have been deposited in SRA, (PRJNA1330141). Data supporting the 5’P-seq results of Figure 4 have been deposited in SRA, (PRJNA1327086). Data supporting the proteomic analysis of Figure 5 and 6 have been deposited in the PRIDE database (reference: 1-20250919-080111-3723971)

## Author Contributions

ADan, AC, QR, JV, AA, JK, CB and AK performed the experiments. ADan, MCC, RM, JV, ADav, CH, JJF, SK, LD and JMD analyzed the data. ADan, RM, JJF, SK, LD and JMD wrote the manuscript.

## Acknowledgments and funding

The Epitranscriptomics and sequencing (EpiRNA-Seq) core facility (University of Lorraine, Nancy, France) is acknowledged. The authors acknowledge S. Koehler and V. Cognat from the Gene expression analysis and the Bioinformatics core facilities of IBMP for their technical help and Audrey Vassord for their technical help, as well as Bastian Welter for technical assistance with metabolite measurements. This work was supported by a grant from the Agence National de la Recherche (ANR-22-CE20-0023-01), by a "New Frontier" grant from the Laboratoires d’Excellences (LABEX)" TULIP (ANR-10-LABX-41), by the CNRS and Université de Perpignan via Domitia. Arnaud Dannfald Salary was financed by the Occitanie Region. This study is set within the framework of the École Universitaire de Recherche TULIP-GS (ANR-18-EURE-0019). S.K. research is funded by the German Research Foundation (DFG) under Germanýs Excellence Strategy – EXC 2048/1 – project 390686111 and within the transregional collaborative research center - TRR341/1 - project 456082119. Mass spectrometry experiments were carried out using the facilities of the Montpellier Proteomics Platform (PPM-PPC, BioCampus Montpellier), member of the Proteomics French Infrastructure (ProFI).

---

**Supplementary Figure S1. Comparison of chemical marks detected outside of the anticodon loop between purified tRNA^Lys^(UUU) (A) and tRNA^Gln^(UUG) (B) in Col0 (WT), *ctu1*, *ctu2* and *trm9* backgrounds**. The y-axis of the histogram represents the ratio of the area under the curve (AUC) measured by liquid chromatography/mass spectrometry (LC-MS/MS) of the modified nucleoside (N*) relative to the corresponding unmodified nucleoside (N).

**Supplementary Figure S2. Analysis by mim-tRNAseq of modifications at position 32, 37 and 58.** Integrative genomics viewer (IGV) screenshot showing examples of sequencing reads (red bars) mapping the tRNA^Lys^(UUU), tRNA^Glu^(UUC) and tRNA^Gln^(UUG). Data from biological triplicates of WT, *ctu1* and *ctu2* mutants are presented for tRNA^Lys^(UUU) while only WT results are shown for the two other tRNAs. Reference nucleotide sequences from position are presented. Two other modifications, m^2^_2_G at position 26 of tRNA^Lys^(UUU), and m^2^G at position 9 of tRNA^Gln^(UUG) are indicated.

**Supplementary Figure S3. Impact of the *ctu2* mutation on global translation.** Polysome profiling of Col-0 (red) and *ctu2* (light blue) using Arabidopsis 15-days-old shoots (A) or roots (B) or 5-days-old whole seedlings (C). The histograms (D to F) show the quantification of low and high polysomes in Col0 (red) and *ctu2* (light blue), measured as the ratio of polysomes to total area (i.e., the area between the 40S, 60S, monosomes and polysomes and the basal lines) in the three situations. The abundance of low polysomes (2-4 ribosomes) is similar between the two genotypes but a significant increase (*P<0.05, T-test n=4) of high polysomes (>4 ribosomes) is observed in the *ctu2* mutant compared to Col0 for 15-days-old shoots (A) and 5-days-old whole seedlings (C), ns: not significant. Error bars represent the standard deviation (SD) of four biological replicates.

