## Supplementary Figures for "Loss of tRNA uridine thiolation affects mRNA translation, protein production, and sulfur-compound metabolism in Arabidopsis"

### Figure S1

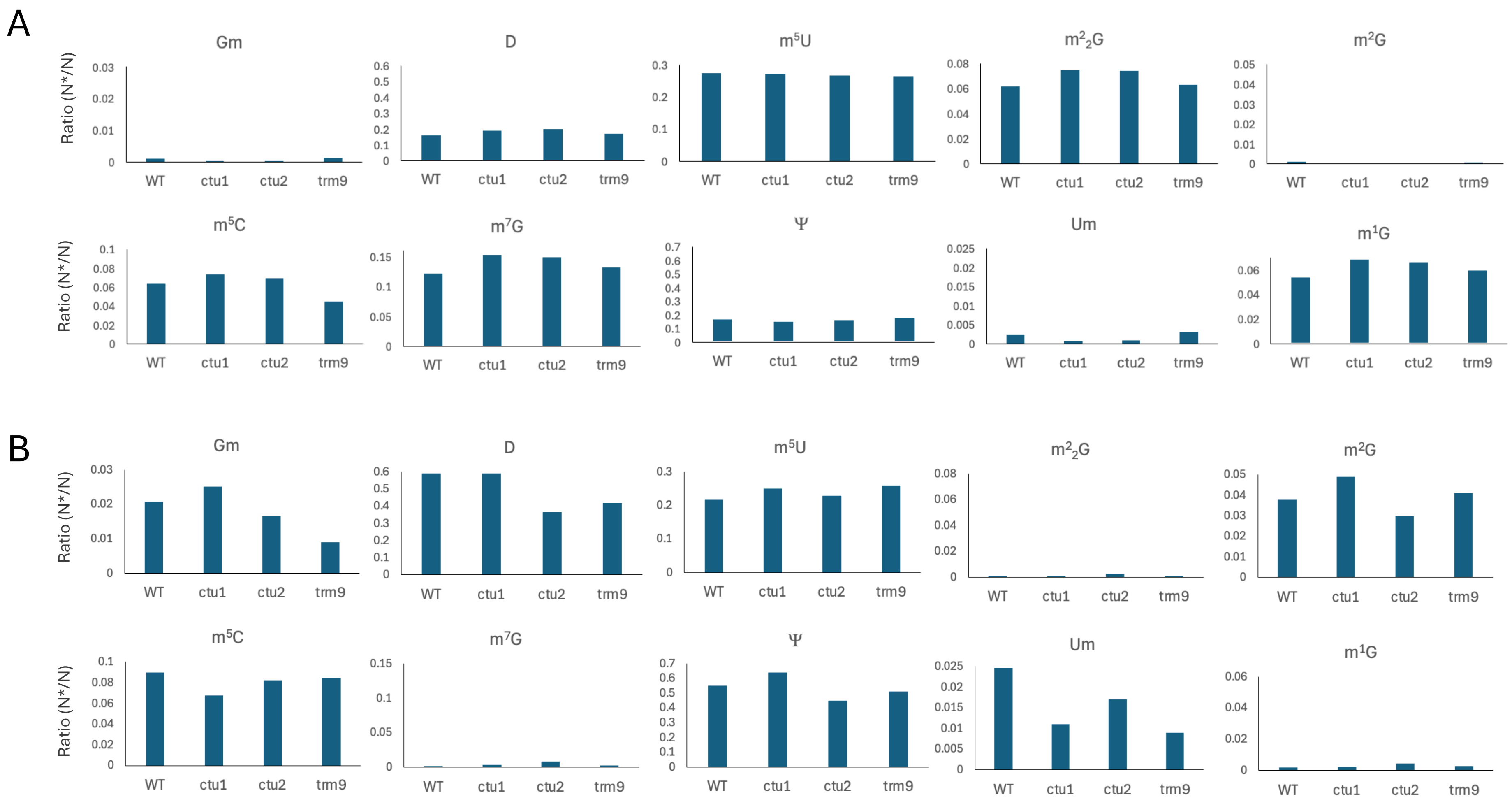


**Supplementary Figure S1.** Comparison of chemical marks detected outside of the anticodon loop between purified tRNA$^{Lys}$(UUU) (A) and tRNA$^{Gln}$(UUG) (B) in Col0 (WT), *ctu1*, *ctu2* and *trm9* backgrounds. The y-axis of the histogram represents the ratio of the area under the curve (AUC) measured by liquid chromatography/mass spectrometry (LC-MS/MS) of the modified nucleoside (N*) relative to the corresponding unmodified nucleoside (N).

### Figure S2

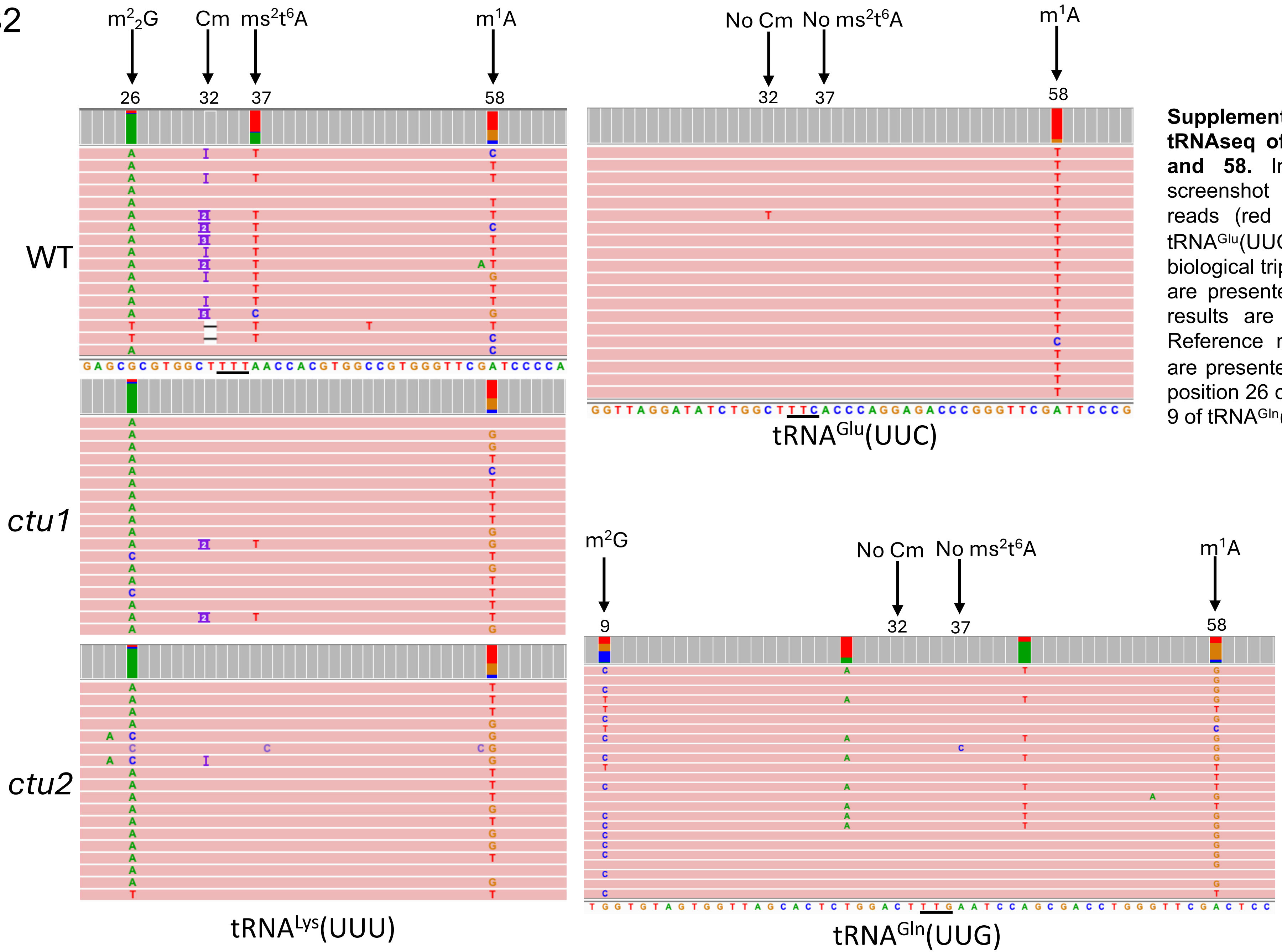


**Supplementary Figure S2. Analysis by mim-tRNAseq of modifications at position 32, 37 and 58.** Integrative genomics viewer (IGV) screenshot showing examples of sequencing reads (red bars) mapping the tRNA$^{Lys}$(UUU), tRNA$^{Glu}$(UUC) and tRNA$^{Gln}$(UUG). Data from biological triplicates of WT, *ctu1* and *ctu2* mutants are presented for tRNA$^{Lys}$(UUU) while only WT results are shown for the two other tRNAs. Reference nucleotide sequences from position are presented. Two other modifications, m$^2_2$G at position 26 of tRNA$^{Lys}$(UUU), and m$^2$G at position 9 of tRNA$^{Gln}$(UUG) are indicated.

### Supplementary Figure S3

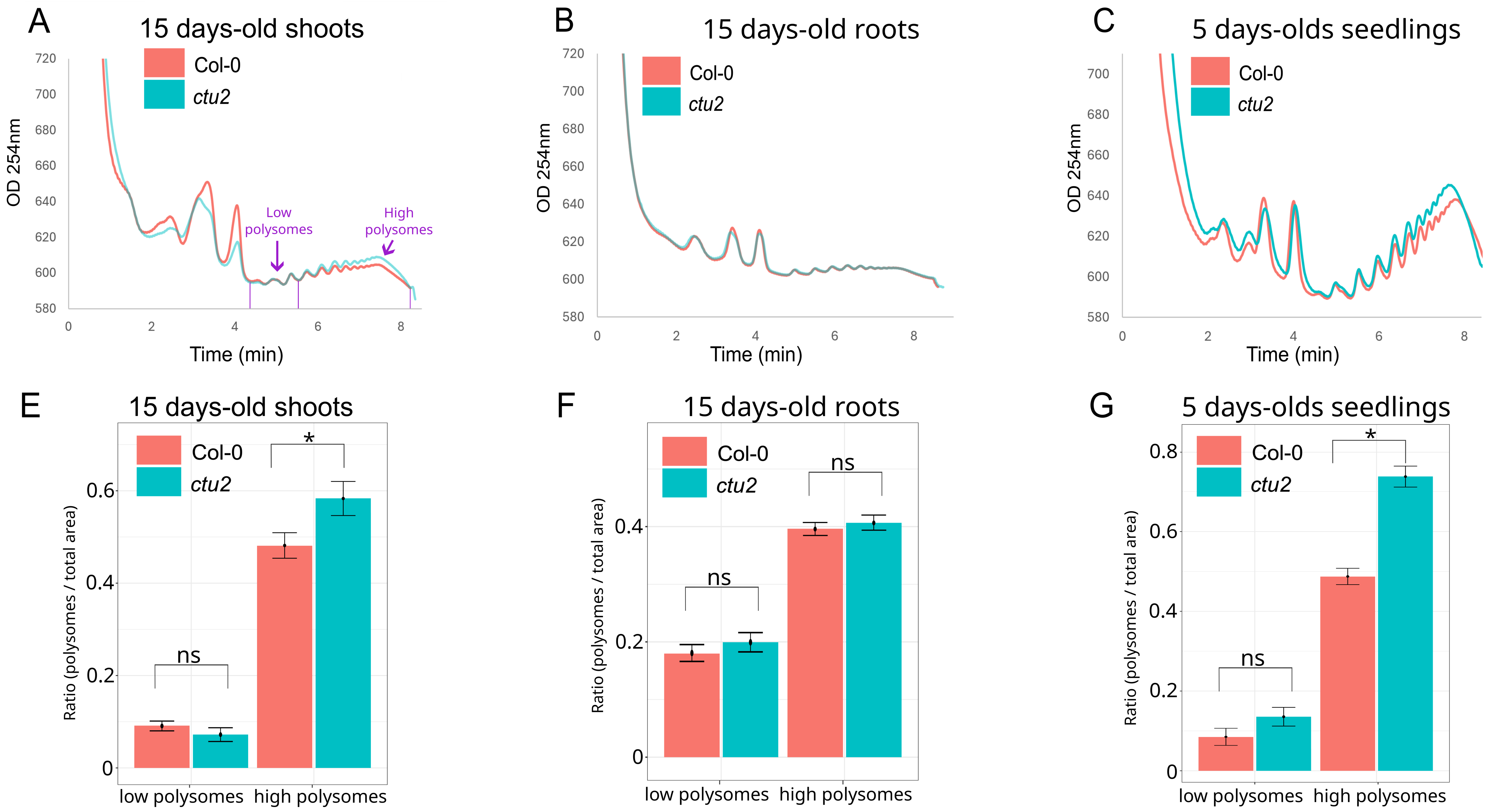


**Supplementary Figure S3**. Impact of the *ctu2* mutation on global translation. Polysome profiling of Col-0 (red) and *ctu2* (light blue) using Arabidopsis 15-days-old shoots (A) or roots (B) or 5-days-old whole seedlings (C). The histograms (D to F) show the quantification of low and high polysomes in WT (Col0, red) and *ctu2* (light blue), measured as the ratio of polysomes to total area (i.e., the area between the 40S, 60S, monosomes and polysomes and the basal lines) in the three situations. The abundance of low polysomes (2-4 ribosomes) is similar between the two genotypes but a significant increase (*P<0.05, T-test, n=4) of high polysomes (>4 ribosomes) is observed in the *ctu2* mutant compared to Col0 for 15-days-old shoots (A) and 5-days-old whole seedlings (C), ns: not significant. Error bars represent the standard deviation (SD) of four biological replicates.
