## Supplementary material for "Loss of tRNA uridine thiolation affects mRNA translation, protein production, and sulfur-compound metabolism in Arabidopsis": Table S4

### Supplementary Table S4

**Table S4A: ESI source parameters for the detection of nucleosides**

| Parameter | Values |
|---|---|
| Nebulizer Gas Flow | 3 L/min |
| Heating Gas Flow | 10 L/min |
| Interface Temperature | 350°C |
| Desolvation Temperature | 602°C |
| DL Temperature | 225°C |
| Heat Block Temperature | 400°C |
| Drying Gas Flow | 3 L/min |
| Interface Voltage | 0.5 kV |
| Focus Voltage | 2 kV |

**Table S4B: ESI source parameters for the detection of nucleosides**

| Nucleoside | Precursor m/z | Products m/z | Target Q1 Pre Bias (V) | Target Collision Energy (V) | Target Q3 Pre Bias (V) | Ret.Time (min) |
|---|---|---|---|---|---|---|
| A | 268 | 136 | -12 | -16 | -15 | 9.0 |
| ac4C | 286.1 | 154 | -10 | -10 | -16 | 11.2 |
| | 286.1 | 112 | -13 | -27 | -12 | |
| acp3U | 346 | 214 | -17 | -16 | -25 | 4.0 |
| | 346 | 197 | -17 | -22 | -22 | |
| | 346 | 168 | -17 | -19 | -19 | |
| Am | 282 | 136 | -13 | -15 | -15 | 11.0 |
| C | 244.1 | 112 | -18 | -13 | -12 | 2.8 |
| Cm | 258.1 | 112 | -18 | -11 | -13 | 7.5 |
| cm5U | 303 | 171 | -15 | -9 | -19 | 7.5 |
| | 303 | 125 | -15 | -24 | -24 | |
| | 303 | 153 | -15 | -16 | -17 | |
| D | 247.1 | 133 | -15 | -7 | -10 | 2.5 |
| | 247.1 | 115 | -16 | -7 | -13 | |
| | 247.1 | 97 | -11 | -19 | -22 | |
| f5C | 272.1 | 140.1 | -14 | -12 | -16 | 8.8 |
| G | 284.1 | 152 | -13 | -12 | -17 | 8.5 |
| Gm | 298.1 | 152 | -11 | -12 | -11 | 10.5 |
| ho5U | 261.2 | 129 | -12 | -10 | -22 | 3.6 |

| Nucleoside | Precursor m/z | Products m/z | Target Q1 Pre Bias (V) | Target Collision Energy (V) | Target Q3 Pre Bias (V) | Ret.Time (min) |
|---|---|---|---|---|---|---|
| I | 269.1 | 137 | -13 | -11 | -15 | 8.3 |
| i6A | 336.15 | 204 | -25 | -17 | -15 | 20.8 |
| | 336.15 | 148 | -19 | -23 | -17 | |
| | 336.15 | 136 | -22 | -33 | -26 | |
| io6A | 352.2 | 220 | -15 | -17 | -25 | 16.8 |
| | 352.2 | 136 | -12 | -35 | -27 | |
| | 352.2 | 118 | -15 | -45 | -28 | |
| m1A | 282.1 | 150.1 | -13 | -18 | -16 | 4.7 |
| m1acp3Y | 360.1 | 228 | -26 | -15 | -26 | 4.6 |
| | 360.1 | 139 | -13 | -31 | -16 | |
| | 360.1 | 270 | -18 | -15 | -30 | |
| m1G | 298.1 | 166 | -14 | -14 | -18 | 10.2 |
| m1I | 283.1 | 151 | -18 | -10 | -17 | 10.5 |
| m2,2G | 312 | 180 | -14 | -14 | -19 | 12.8 |
| m27G | 312.1 | 180 | -14 | -16 | -13 | 9.6 |
| m2A | 282.1 | 150 | -12 | -17 | -17 | 9.6 |
| m2G | 298 | 166 | -13 | -13 | -18 | 10.8 |
| m3C | 258.1 | 126 | -12 | -14 | -14 | 3.7 |
| m3U | 259 | 127 | -12 | -11 | -14 | 9.9 |
| m5C | 258 | 126 | -12 | -12 | -14 | 4.7 |
| m5Cm | 272.1 | 126.2 | -14 | -13 | -15 | 9.0 |
| m5U | 259 | 127 | -30 | -11 | -14 | 8.8 |
| m5Um | 273.1 | 127 | -14 | -11 | -15 | 12.3 |
| m7G | 298.3 | 166 | -10 | -15 | -18 | 7.3 |
| mcm5s2U | 333.1 | 201 | -10 | -9 | -23 | 14.2 |
| | 333.1 | 169 | -15 | -18 | -12 | |
| | 333.1 | 141 | -12 | -28 | -15 | |
| mcm5U | 317.1 | 185 | -15 | -9 | -10 | 10.8 |
| | 317.1 | 153 | -14 | -18 | -16 | |
| | 317.1 | 125 | -15 | -28 | -21 | |
| mcm5Um | 331 | 185 | -17 | -10 | -21 | 14.2 |
| | 331 | 125 | -17 | -26 | -24 | |
| | 331 | 153 | -16 | -19 | -30 | |
| mnm5s2U | 304.1 | 172.2 | -11 | -12 | -20 | 5.3 |
| | 304.1 | 140.9 | -11 | -17 | -11 | |
| | 304.1 | 255.2 | -15 | -11 | -20 | |
| | 304.1 | 273.1 | -15 | -10 | -15 | |
| ms2t6A | 458.9 | 327.1 | -13 | -14 | -18 | 18.0 |
| ncm5s2U | 318 | 186 | -11 | -10 | -14 | 8.6 |
| | 318 | 169 | -11 | -17 | -13 | |
| | 318 | 141 | -20 | -28 | -29 | |
| ncm5U | 302.1 | 170 | -15 | -10 | -18 | 4.4 |

| Nucleoside | Precursor m/z | Products m/z | Target Q1 Pre Bias (V) | Target Collision Energy (V) | Target Q3 Pre Bias (V) | Ret.Time (min) |
|---|---|---|---|---|---|---|
| | 302.1 | 153 | -14 | -17 | -11 | |
| | 302.1 | 125 | -13 | -29 | -28 | |
| Pseudouridine (ψ) | 245.05 | 209 | -21 | -9 | -24 | 2.7 |
| | 245.05 | 179 | -19 | -11 | -19 | |
| | 245.05 | 155 | -11 | -13 | -26 | |
| R-mchm5U | 333 | 183 | -17 | -16 | -21 | 9.0 |
| | 333 | 201 | -17 | -9 | -23 | |
| s2C | 260.3 | 128 | -12 | -11 | -14 | 6.1 |
| | 260.3 | 111 | -13 | -37 | -23 | |
| S-mchm5U | 333 | 183 | -17 | -16 | -21 | 8.2 |
| | 333 | 201 | -17 | -9 | -23 | |
| t6A | 413.1 | 281 | -21 | -13 | -21 | 16.5 |
| | 413.1 | 136 | -15 | -31 | -29 | |
| | 413.1 | 162 | -12 | -25 | -19 | |
| U | 245.1 | 113 | -19 | -9 | -20 | 5.0 |
| Um | 259.1 | 113 | -12 | -9 | -13 | 9.7 |
